# Expression of m^6^A RNA methylation markers in the hypothalamus of Atlantic salmon

**DOI:** 10.1101/2022.09.08.507106

**Authors:** Ehsan Pashay Ahi, Morgane Frapin, Mikaela Hukkanen, Craig R. Primmer

## Abstract

Methylation at the N6-position of adenosine, m^6^A, is the most abundant mRNA modification in eukaryotes. It is a highly conserved universal regulatory mechanism controlling gene expression in a myriad of biological processes. The role of m^6^A methylation in sexual maturation, however, has remained largely unexplored. While the maturation process is known to be affected by many genetic and environmental factors, the molecular mechanisms causing variation in the timing of maturation are still poorly understood. Hence, investigation of whether a widespread mechanism like m^6^A methylation could be involved in controlling of the maturation timing is warranted. In Atlantic salmon (*Salmo salar*), two genes associated with the age at maturity in human, *vgll3* and *six6*, have been shown to play an important role in maturation timing. In this study, we investigated the expression of 16 genes involved in the regulation of m^6^A RNA methylation in the hypothalamus of Atlantic salmon with different homozygous combinations of *late* (L) and *early* (E) alleles for *vgll3* and *six6* genes. We found differential expression of *ythdf2*.*2* which encodes an m^6^A modification reader and promotes mRNA degradation. Its expression was higher in *six6*LL* compared to other genotypes as well as immature males compared to matures. In addition, we found that the expression levels of genes coding for an eraser, *alkbh5*, and for a reader, *ythdf1*, were higher in the hypothalamus of females than in males across all the different genotypes studied. Our results indicate a potential role of the m^6^A methylation process in sexual maturation of Atlantic salmon, and therefore, provide the first evidence for such regulatory mechanism in the hypothalamus of any vertebrate. Investigation of additional vertebrate species is warranted in order to determine the generality of these findings.

## Introduction

Several post-transcriptional mechanisms allow cells to regulate gene expression by acting at the level of transcription and/or translation, such as microRNAs, alternative splicing and RNA methylation (Cai et al., 2009; Frye and Blanco, 2016; Singh and Ahi, 2022). These mechanisms are essential to produce a vast cellular diversity from cells with an identical genome but also to adapt to changing environments at the organismal level. Methylation at the N6 position of adenosine (so called N6-Methyladenosine or m^6^A) is the most abundant and highly conserved mRNA modification in eukaryotes, which has recently emerged as a universal regulatory mechanism controlling gene expression in myriad of biological processes (Zaccara et al., 2019). The N6-Methyladenosine, m^6^A, controls the fate of mRNAs within cells by acting on processes such as mRNA stability, splicing and transport. These modifications are added by the m^6^A methyltransferase complex, which includes proteins called writers (e.g. Mettl3, Mett14 and Wtap), and can be removed by demethylases called erasers (e.g. Fto and Alkbh5) (Zaccara et al., 2019). The RNA m^6^A modifications are recognized by proteins called readers (e.g. Ythdf1/2/3 and Ythdc1/2) that guide methylated mRNA towards specific fates such as degradation, stabilization, transportation, and promotion or inhibition of translation (Liao et al., 2018).

The m^6^A RNA modification has been recently found to be crucially involved in reproduction, and the tip of the iceberg has just started to be revealed by findings on the essential role of m^6^A modification during gametogenesis (Lasman et al., 2020; Xia et al., 2018). However, there is no knowledge on the potential role of m^6^A modification upstream of initiating and orchestrating sexual maturation along hypothalamus-pituitary-gonadal (HPG) axis, and hence its possible role in contributing to variation in maturation timing. The complex molecular mechanisms causing variation in maturation timing are generally poorly understood (Howard and Dunkel, 2019; Leka-Emiri et al., 2017; Mobley et al., 2021), which makes it even more tempting to investigate whether a universal mechanism such as m^6^A RNA modification may contribute to maturation timing variation.

Age at maturity, a critical trait directly determined by mechanisms controlling maturation timing, affects fitness traits such as survival and reproductive success (Mobley et al., 2021). Particularly when it comes to an economically important and ecologically vulnerable fish species such as Atlantic salmon (*Salmo salar*), understanding the underlying biology of age at maturity is of paramount importance. Interestingly, two genes that have been associated with age at maturity in human, *VGLL3* and *SIX6* (Cousminer et al., 2016; Perry et al., 2014), also play a major role in pubertal timing of Atlantic salmon (Barson et al., 2015; Sinclair-Waters et al., 2020). The gene *vgll3* (the vestigial-like family member 3 gene) is strongly associated with maturation timing in both sexes of wild Atlantic salmon but also exhibits sex-specific maturation effects (Barson et al., 2015; Czorlich et al., 2018). This association between *vgll3* genotype and maturation probability has been validated in one year-old male parr in common garden settings (Debes et al., 2021; Sinclair-Waters et al., 2021; Verta et al., 2020). In our recent studies, we also showed that *vgll3* strongly affects the expression of reproductive axis genes in one year-old male Atlantic salmon (Ahi et al., 2022b), and its regulatory effects on transcription of the gonadotropin encoding genes (*fshb* and *lhb*) are predicted to be mediated by the Hippo signaling pathway (Ahi et al., 2022a). The molecular mechanism by which *six6* (sine oculis homeobox 6) regulates pubertal timing is not known. However, it seems that *six6* transcriptional regulation is not linked to *vgll3* function and it is independent of Hippo signaling (Kurko et al., 2020).

In this study, we investigate the expression of 16 genes that have been found to code proteins associated with the regulation of m^6^A modifications including 3 writers; *mettl3, mettl14*, and *wtap*, 4 erasers; *alkbh5-1, alkbh5-2, fto-1* and *fto-2*, and 9 readers; *ythdc1-1, ythdc1-2, ythdc2, ythdf1-1, ythdf1-2, ythdf1-3, ythdf2-1, ythdf2-2* and *ythdf3*. We compare their mRNA expression between different homozygous combinations of *late* (L) and *early* (E) alleles for *vgll3* and *six6* genes in the hypothalamus of both sexes of one year old Atlantic salmon. In males, we also compared the expression of m^6^A methylation markers between immature and mature individuals. To our knowledge, this is the first study addressing differences in the expression of m^6^A methylation markers in respect to sexual maturation in the hypothalamus. These findings provide the foundation for future investigation of the role of m^6^A methylation in HPG axis control of sexual maturation in fish and other vertebrates.

## Materials and methods

### Fish rearing, genotyping and tissue sampling

The Atlantic salmon used in this study were created using parental individuals from a 1^st^ generation hatchery broodstock originating from the Iijoki population that is maintained at the Natural Resources Institute (Luke) Taivalksoki hatchery in northern Finland. Unrelated parents were chosen from broodstock individuals that had earlier been genotyped for 177 SNPs on Ion Torrent or Illumina (Miseq or Next-Seq) sequencing platforms as outlined in Aykanat et al., 2016. These SNPs included SNPs linked to the *vgll3* and *six6* genes that were earlier shown to be associated with age at maturity in salmon (Barson et al., 2015). We selected parents that were heterozygotes for both *vgll3* and *six6* (*vgll3*EL* and *six6*EL*) in order for full-sib families to contain offspring with all *vgll3* - *six6* genotype combinations. We avoided crossing closely related individuals (those with grandparents in common) by using SNP-based pedigree reconstruction as in Debes et al., 2021. From this point onwards, four character genotypes will be used to describe an individual’s genotype at the focal loci, *vgll3* and *six6*. The first two characters indicate the genotype at the *vgll3* locus and the last two characters indicate the genotype at the *six6* locus. The locus is indicated in subscript text after the genotype. In order to minimize unwanted variation, all individuals in this study originate from a single full-sib family. For simplicity, only the four homozygous genotype combinations (*vgll3*EE* or *vgll3*LL* and *six6*EE* or *six6*LL*) were examined. This enabled the assessment of the expression patterns of all the possible homozygous *vgll3* and *six6* genotypes within an otherwise similar genetic background. Other rearing, tagging and genotyping details are as described in Sinclair-Waters et al., 2021.

The fish were euthanized approximately one-year post-fertilization during the spawning season in November with an overdose of the anesthetic buffered tricaine methane sulfonate (MS-222) and dissected, and sex and maturation status were determined visually by observing the presence of female or male gonads as outlined in Verta et al., 2020. The maturation status of males was classified as immature (no phenotypic signs of gonad maturation) and mature (large gonads leaking milt). The mature or immature status of male salmon was determined by respectively the presence or the absence of sperm leakage. At this age all the females were immature. Whole hypothalami of salmon with the genotypes of interest were dissected and snap-frozen in liquid nitrogen before being stored at −80 °C.

### RNA extraction and cDNA synthesis

RNA was isolated using NucleoSpin RNA kit (Macherey-Nagel GmbH & Co. KG). The hypothalami were transferred to tubes with 1.4 mm ceramic beads (Omni International), Buffer RA1 and DDT (350ul RA1 and 3,5ul DDT 1M) and homogenized using Bead Ruptor Elite (Omni International) with tissue specific program (4m/s, 3×20s). Remaining steps of RNA isolation were conducted as in the manufacturer’s protocol. RNA was eluted in 40 µl of nuclease free water. Quality and concentration of the samples were measured with NanoDrop ND-1000. The extracted RNA (400ng for females, 500ng for males per sample) was subsequently reverse-transcribed to cDNA using iScript cDNA Synthesis Kit (Bio-Rad Laboratories, Inc.)

### Primer design and qPCR

We used paralogue-specific gene sequences obtained from the recently annotated *Salmo salar* genome in the Ensembl database, http://www.ensembl.org. The paralogue gene sequences were aligned using CLC Genomic Workbench (CLC Bio, Denmark) in order to identify paralogue specific regions for designing the primers (Ahi et al., 2014). The steps of primer designing are as described in Ahi et al., 2019, using two online tools: Primer Express 3.0 (Applied Biosystems, CA, USA) and OligoAnalyzer 3.1 (Integrated DNA Technology) (Supplementary data). The qPCR reactions were prepared as described in Ahi et al., 2018, using PowerUp SYBR Green Master Mix (Thermo Fischer Scientific), and performed on a Bio-Rad CFX96 Touch Real Time PCR Detection system (Bio-Rad, Hercules, CA, USA). The details of the qPCR program and calculation of primer efficiencies are described in Ahi et al., 2019.

### Data analysis

The Cq values of the target genes were normalized with the geometric mean of the Cq values of two references genes, *elf1a* and *hprt1*, following the formula ΔCq _target_ = Cq _target_ – Cq _reference_. These ΔCq _target_ values have been adjusted for qPCR batch effect using ComBat (v3) (Johnson et al., 2007) from sva R package (Surrogate Variable Analysis, v3.40.0) (Leek et al., 2012). The following calculations have been made on the qPCR batch adjusted values. For each gene, a biological replicate with the lowest expression level across all the used in each comparison (calibrator sample) was selected to calculate ΔΔCq values (ΔCq _target_ – ΔCq _calibrator_). The relative expression quantities (RQ values) were calculated as 2^−ΔΔCq^, and their fold changes (logarithmic values of RQs) were used for statistical analysis (Pfaffl, 2001). The effects of genotype and maturity status were modeled separately: the between-genotype effects were examined within maturity stages, whereas the between-maturity stages were examined within each genotype. An ANOVA (Analysis of variance) test was applied for the analysis between genotypes. Benjamini-Hochberg method (Benjamini & Hochberg 1995) was used for multiple-comparison correction. In cases where the variable of interest (genotype or maturity) explained variation significantly (p<0.10) after correcting for multiple comparisons, further post-hoc tests were conducted. Post-hoc tests were calculated using Tukey’s Honest Significant Difference test. Statistical analyses were conducted using R software (version 4.1.1).

## Results

### Expression levels of m^6^A methylation markers in hypothalamus of Atlantic salmon

We first assessed the overall expression levels of genes encoding for proteins involved in the regulation of m6A methylation in males (maturity status separately) and in females (Fig. 1). Among the writers, *wtap* had the highest expression level (lowest dCq values), whereas *mettl3* displayed the lowest expression level (highest dCq values) (Fig. 1A). The paralogous genes of the two erasers showed different expression levels as well, i.e. *alkbh5-1* > *alkhb5-2* and *fto-1* > *fto-2* (Fig. 1B). Similarly, the paralogous genes of three readers showed differences in their expression level, i.e. *ythdc1-2* > *ythdc1-1, ythdf1-3* > *ythdf1-2* > *ythdf1-1*, and *ythdf2-1* > *ythdf2-2* (Fig. 1C). However, the expression level difference between *ythdf2-1* and *ythdf2-2* appeared to be minor compared to the differences between the paralogs of the other genes. Among the readers, *ythdf1-3* had the highest expression level, whereas *ythdc2* exhibited the lowest expression level (Fig. 1C). In general, the sex and maturity status did not seem to have any effect on expression level differences between the paralogs.

**Figure 1:**
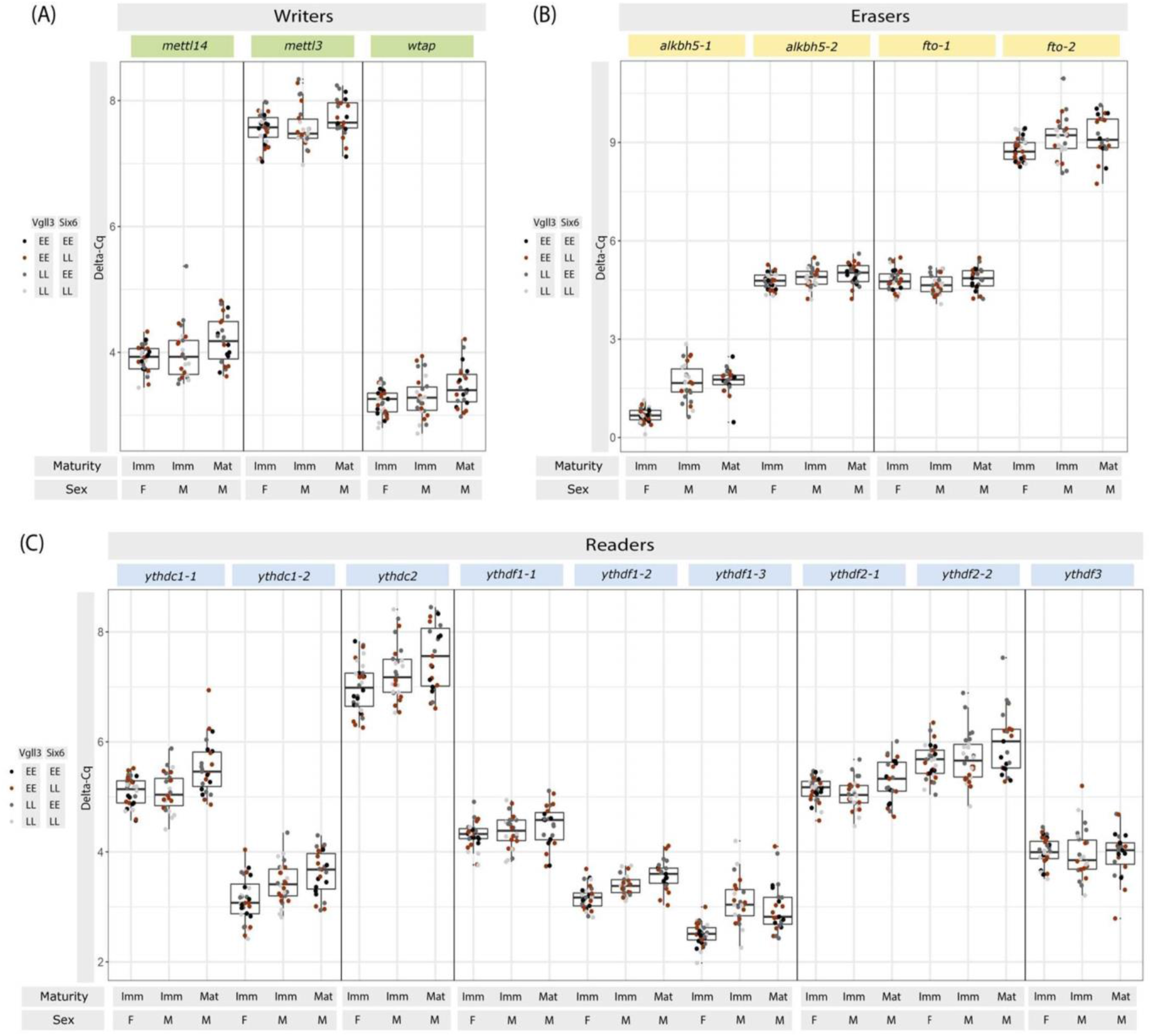
Overall mRNA expression level of the m^6^A RNA modification regulators. qPCR batch adjusted Delta-Cq of the genes coding for writers (A), erasers (B) and readers (C). The boxplots represent the median, first and third quartiles of all the genotypes within the group. (F: Females, M: Males, Imm: Immature, Mat: Mature).

### Expression differences of m^6^A methylation markers between the genotypes

Next, we explored the expression differences of the m6A methylation regulators between the genotypes within each sex and maturity status. In immature males, we found expression differences between the genotypes for only one paralogue of a reader gene, *ythdf2-2*, which displayed higher expression in the genotype combinations with homozygous *late* allele of *six6* (*six6*LL*) (Fig. 2). We did not observe any *vgll3* or *six6* genotype-specific expression difference in the hypothalamus of either mature males or immature females (Fig. 3 and 4).

**Figure 2:**
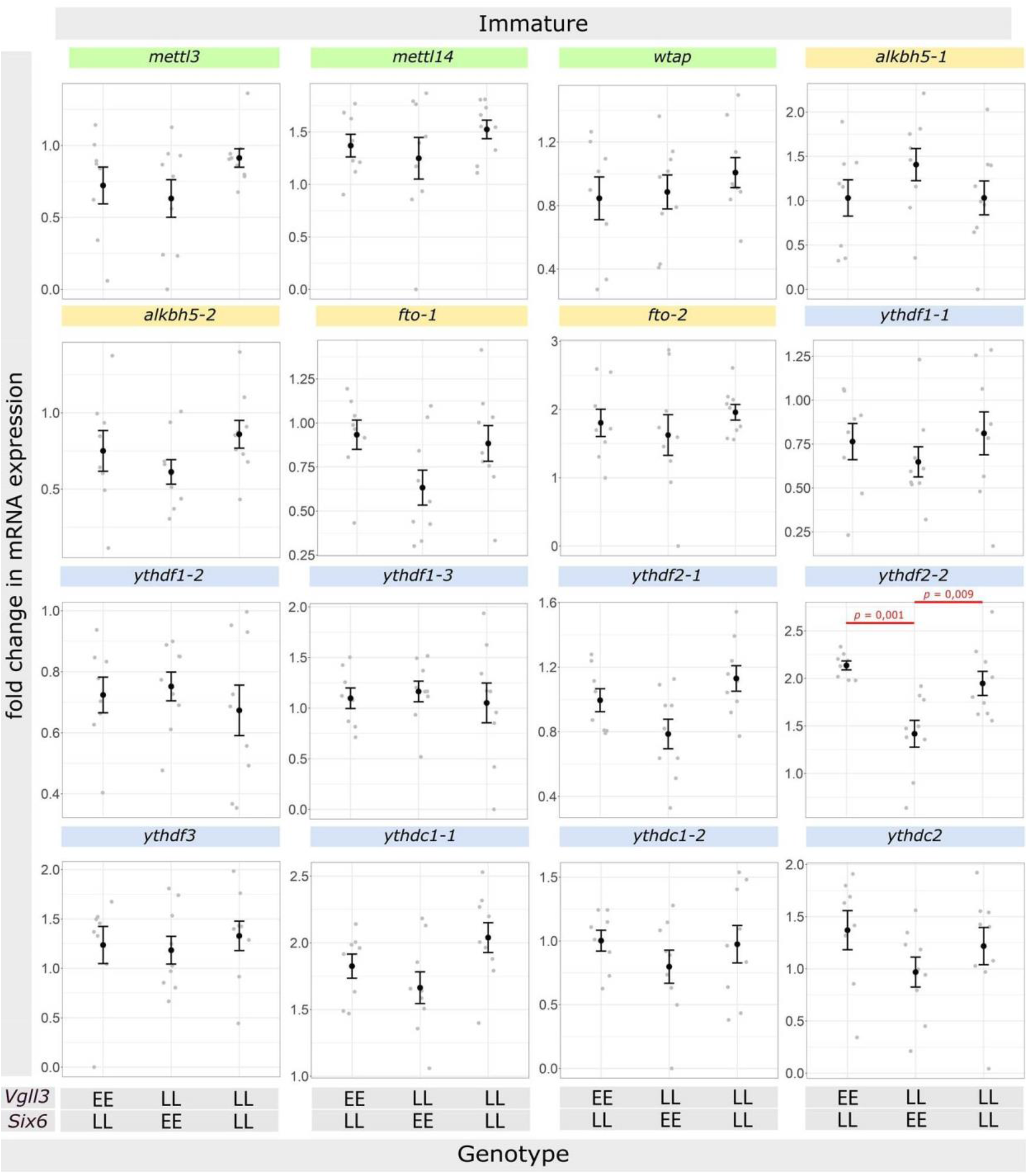
Comparison of the m^6^A RNA modification regulators mRNA expression level between the genotypes in immature males. For each gene, values were Log2 Fold-Change of the mRNA expression for each sample (grey dot) and mean ± SEM (black dot and bar). The mRNA level expression differences between the genotypes were analyzed with ANOVA followed by Tukey HSD (Honest Significant Difference) post-hoc tests.

**Figure 3:**
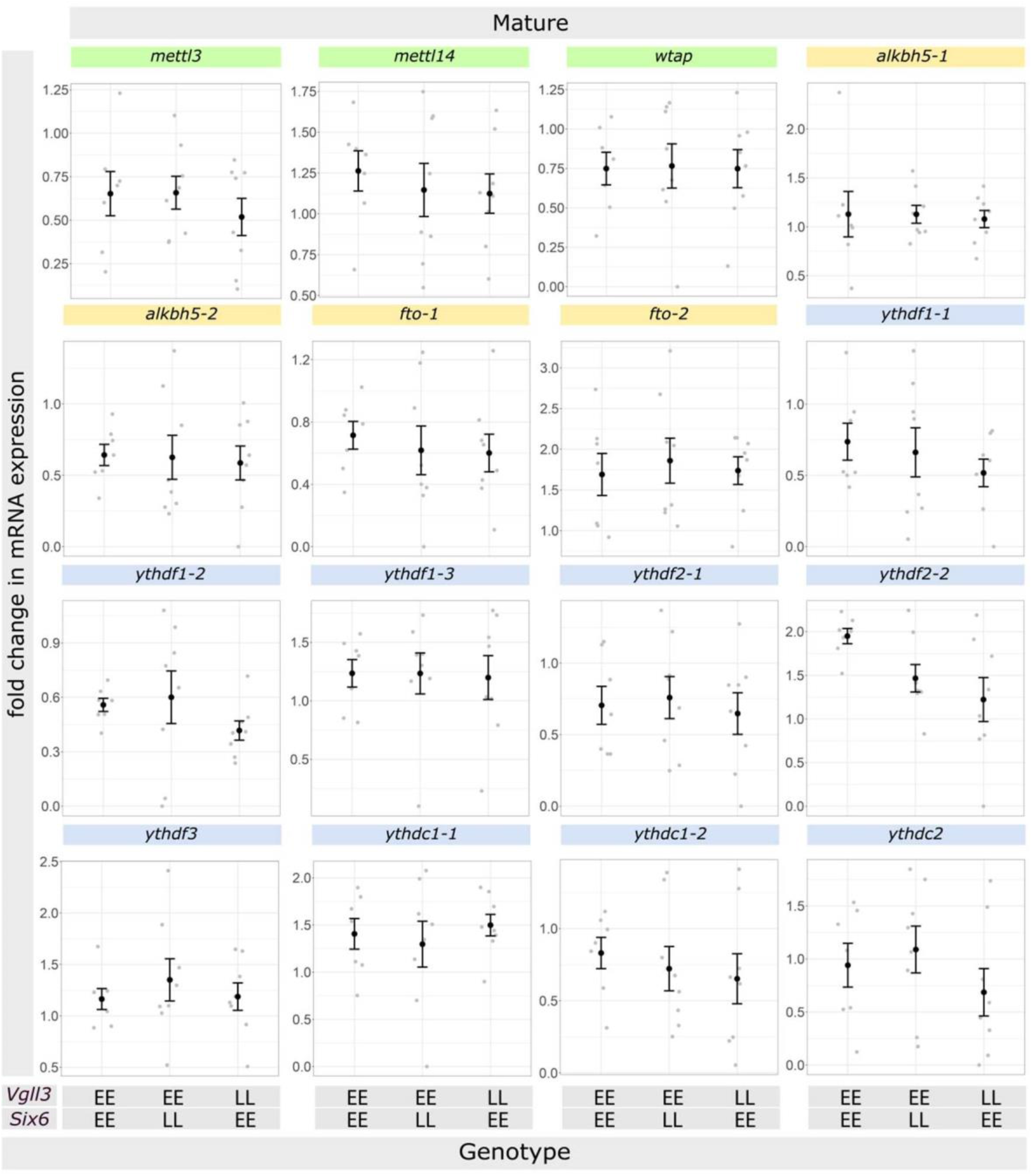
Comparison of the m^6^A RNA modification regulators mRNA expression level between the genotypes in mature males. For each gene, values were Log2 Fold-Change of the mRNA expression for each sample (grey dot) and mean ± SEM (black dot and bar). The mRNA level expression differences between the genotypes were analyzed with ANOVA followed by Tukey HSD (Honest Significant Difference) post-hoc tests.

**Figure 4:**
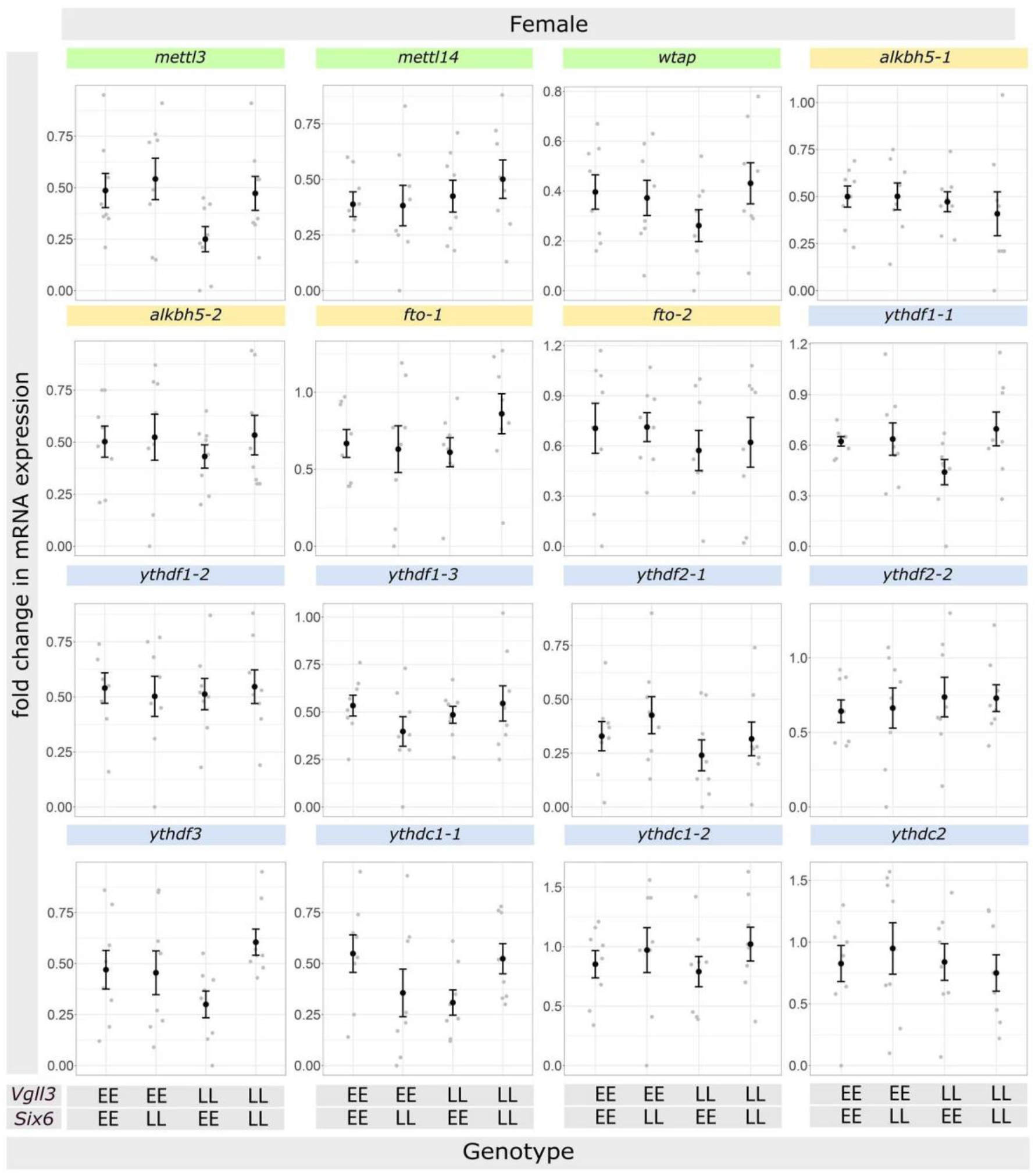
Comparison of the m^6^A RNA modification regulators mRNA expression level between the genotypes in females. For each gene, values were Log2 Fold-Change of the mRNA expression for each sample (grey dot) and mean ± SEM (black dot and bar). The mRNA level expression differences between the genotypes were analyzed with ANOVA followed by Tukey HSD (Honest Significant Difference) post-hoc tests.

### Expression differences of m^6^A methylation markers between the maturity status in males

The comparison between maturity stages was only possible within two genotype combinations *(vgll3*EE/six6*LL* and *vgll3*LL/six6*EE)*, as other genotype combinations did not have both immature and mature males. Within *vgll3*EE/six6*LL* individuals, we observed higher expression of *ythdf2-2* in the hypothalami of immature males compared to mature males (Fig. 5). Whereas, within *vgll3*LL/six6*EE* genotype, we found a paralog of another reader gene, *ythdf1-2*, showing similar expression pattern with higher expression in the hypothalami of immature males compared to mature males (Fig. 6).

**Figure 5:**
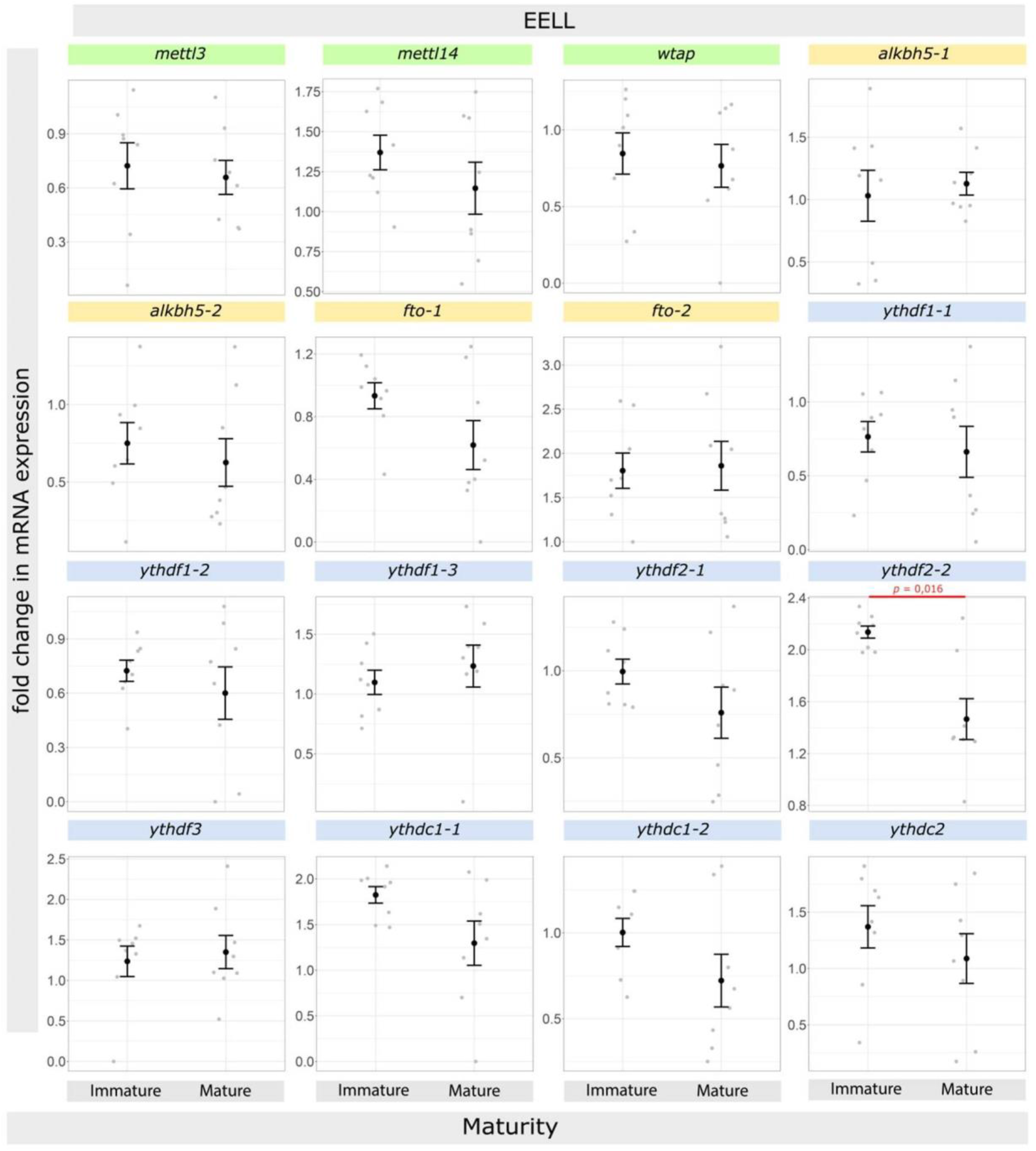
Comparison of the m^6^A RNA modification regulators mRNA expression level between maturity stages in individuals with EE LL genotypes. For each gene, values were Log2 Fold-Change of the mRNA expression for each sample (grey dot) and mean ± SEM (black dot and bar). The mRNA level expression differences between the genotypes were analyzed with ANOVA followed by Tukey HSD (Honest Significant Difference) post-hoc tests.

**Figure 6:**
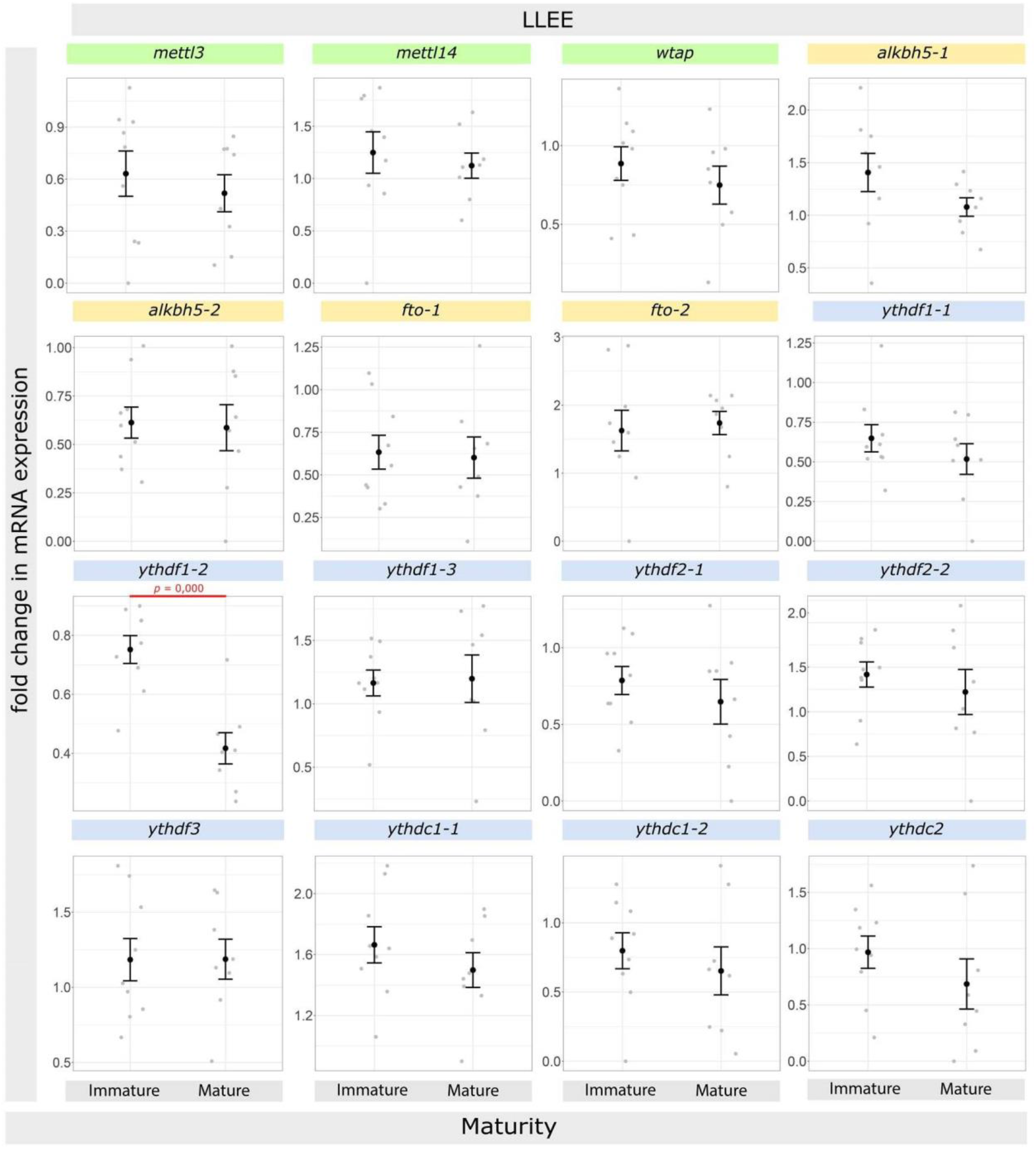
Comparison of the m^6^A RNA modification regulators mRNA expression level between maturity stages in individuals with LL EE genotypes. For each gene, values were Log2 Fold-Change of the mRNA expression for each sample (grey dot) and mean ± SEM (black dot and bar). The mRNA level expression differences between the genotypes were analyzed with ANOVA followed by Tukey HSD (Honest Significant Difference) post-hoc tests.

### Expression differences of m^6^A methylation markers between the sexes

Finally, we compared the expression levels of the m^6^A methylation regulators between sexes within the genotypes classes containing both immature males and females (Fig. 7). We found a paralogous gene of an eraser *alkbh5-1* and a paralogous gene of a reader, *ythdf1-3*, to be differentially expressed between the males and females in all the genotypes. Interestingly, the direction of these expression pattern differences was always the same for both genes; showing higher expression levels in the hypothalamus of females than males. Also, another paralogous gene of the same reader, *ythdf1-2*, showed similar tendency of higher expression in females but the differences were not statistically significant. These findings indicate sex-specific, but not genotype-specific, expression level differences for both of these genes in the hypothalamus of Atlantic salmon.

**Figure 7:**
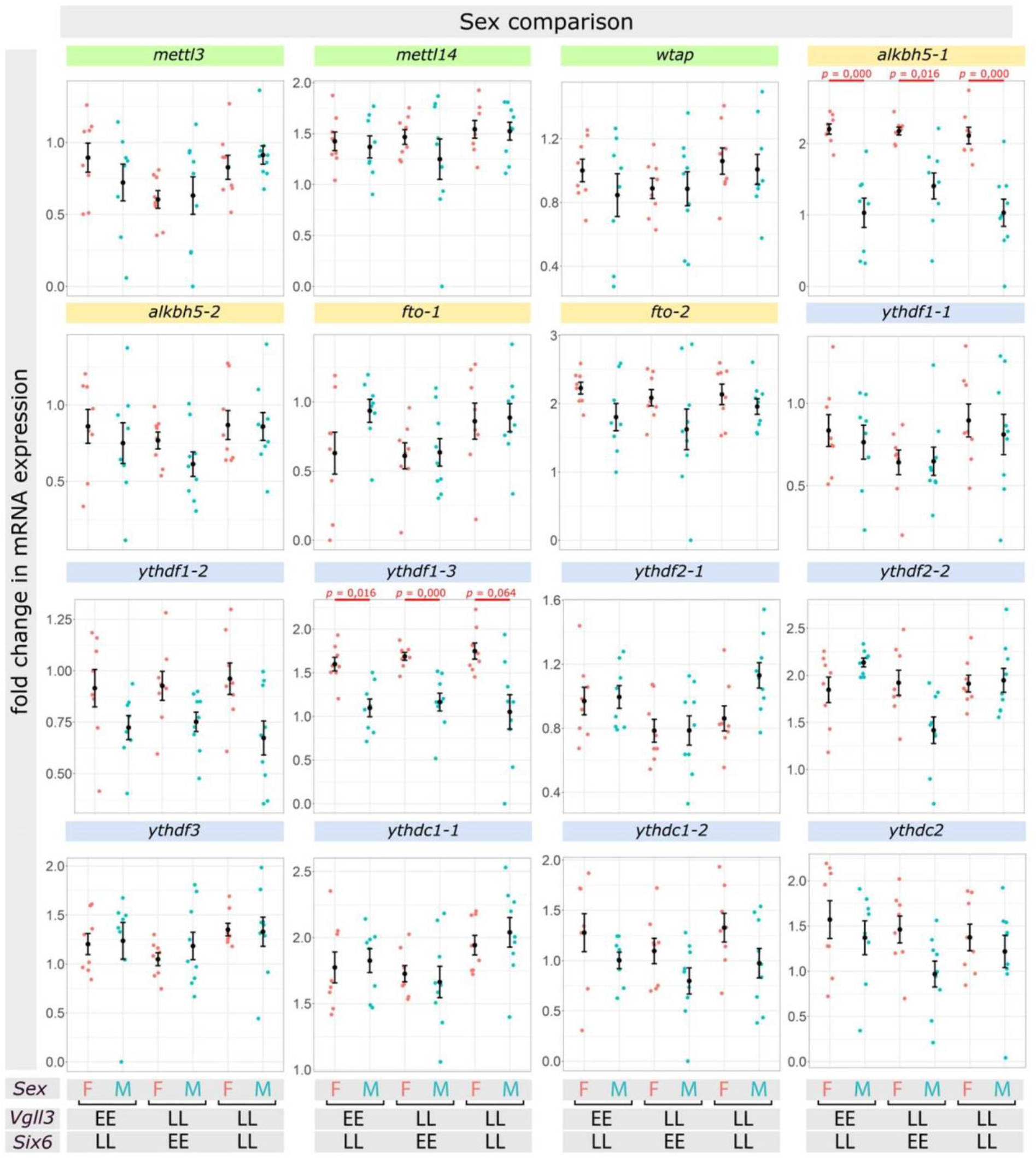
Comparison of the m^6^A RNA modification regulators mRNA expression level between sexes. For each gene, values were Log2 Fold-Change of the mRNA expression for each sample (red and blue dots) and mean ± SEM (black dot and bar). The mRNA level expression differences between the genotypes were analyzed with ANOVA followed by Tukey HSD (Honest Significant Difference) post-hoc tests.

## Discussion

The expression of the m^6^A RNA modification regulators in hypothalamus has never been the focus of research in fish reproduction to date. This is somewhat surprising, not only because the hypothalamus is an important upstream organ regulating sexual maturation (Kah et al., 1993), but also due to the widespread roles of m^6^A RNA methylation, as the most abundant RNA modification in eukaryotic cells, in regulation of numerous biological processes (Zaccara et al., 2019). Particularly, m^6^A RNA modification has been described as an essential mechanism in gonadal development and fertility (Mu et al., 2022). However, in the downstream sex organs, i.e. gonads, the role and expression of some of the m^6^A RNA modification markers have been recently studied in fish (L. Wang et al., 2020; Xia et al., 2018; Zhao et al., 2021). Thus, in this study, we first set out to investigate, for the first time, whether the m^6^A RNA modification regulators are expressed in hypothalamus of a fish species (Atlantic salmon). We found that at least 16 genes encoding the m^6^A RNA modification regulators (writers, erasers and readers) are expressed in the hypothalamus but in variable levels (Fig. 1). Importantly, it appeared that even the paralogs of the same eraser and reader genes can have very variable expression levels in the hypothalamus indicating potential differences in the functional importance of these genes at paralogue level.

The most important finding in this study was the differential expression of *ythdf2-2* in the hypothalamus of male Atlantic salmon (Fig. 2), i.e. with reduced expression in mature compared to immature individuals with the *vgll3*EE/six6*LL* genotype combination, as well as reduced expression in *six6*EE* genotype within immature groups. In Atlantic salmon, there are two paralogous genes (*ythdf2-1* and *ythdf2-2*) for *Ythdf2*, encoding a member of YT521-B homology domain family (YTHDF) proteins, which acts as a reader of m^6^A and its binding to m^6^A-containing RNA leads to degradation of the RNA (Du et al., 2016). In mice, Ythdf2 deficiency results in female infertility and has pivotal role in maternal RNA degradation during oocyte maturation (Ivanova et al., 2017). Another study in zebrafish also reported the pivotal role of *ythdf2* in the maternal-to-zygotic transition by orchestrating the clearance of almost one-third of maternal mRNAs in zygote (Zhao et al., 2017). More recent studies in mice have revealed that Ythdf2-mediated mRNA degradation on m^6^A-modified target transcripts is required for spermatogenesis and fertility (Qi et al., 2022; Zhao et al., 2021). However, no study has investigated the potential role of *Ythdf2* upstream of HPG axis (i.e. hypothalamus) during sexual maturation. In mice, Ythdf2 has been shown to be essential during embryonic neural development by promoting m^6^A-dependent degradation of genes related to neuron differentiation (Li et al., 2018). In chicken hypothalamus, RNA m^6^A modification seems to be involved in the regulation of circadian rhythms under stressful conditions, and the expression m^6^A methylation markers including *Ythdf2* show correlated oscillation with the clock genes (Y. Yang et al., 2022a). Studies in chickens and rats have shown that the level of expression of *Ythdf2* is modified by the environment (i.e. light exposure and maternal diet during gestation) in the hypothalamus (Frapin et al., 2020; Y. Yang et al., 2022b). Interestingly, even though the direct hypothalamic role of *Ythdf2* during sexual maturation has not been explored yet, a decrease in the level of m^6^A methylation by increased hypothalamic expression of *Fto* (a m6A eraser) has been shown to cause early onset of puberty in female rat (X. Yang et al., 2022). The opposite effect of delayed puberty was also obtained by knockdown of *Fto* (X. Yang et al., 2022). The result of the latter study on female rat together with our findings about *ythdf2* indicate that m^6^A modification might be involved in mediated sexual maturation in vertebrates by fine tuning the hypothalamic mRNA levels either through increased expression of a m^6^A eraser (e.g. *Fto* in rat) or reduced expression of m^6^A reader (*ythdf2* in Atlantic salmon).

The genotype dependent expression difference of *ythdf2* was also observed within immature individuals (Fig. 5). This expression difference was more likely to be linked to *six6* genotype than vgll3 genotype, since no difference was observed between *vgll3*LL* and *vgll3*EE* when *six6* genotype remained the same (i.e. it was fixed on *six6*LL*). In respect to *six6* genotype, however, *ythdf2* appeared to have higher expression in *six6*EE* than *six6*LL*. This can be functionally explained, as mentioned above, by the fact that that puberty may favor reduced expression of *ythdf2* and increased level of its mRNA targets in hypothalamus and since *six6*EE* are more prone to enter puberty than *six6*LL*. But this does not explain why a similar pattern was not observed between *vgll3*LL* and *vgll3*EE*, unless there might be an unknown regulatory link only between *six6* and *ythdf2* and not between *vgll3* and *ythdf2*. In mammals, it has been shown that *six6* is involved in development of hypothalamic GnRH neurons, which are essential for the onset of puberty (Pandolfi et al., 2019). Moreover, Ythdf2-mediated mRNA clearance is also demonstrated to be important for neuron maturation in the developing forebrain of mice (Li et al., 2018). However, deeper understanding of the connection between *six6* genotypes and *ythdf2* expression requires further investigations.

Since no direct regulatory link was identified between *six6* and *ythdf2*, one plausible scenario is an indirect hierarchical regulatory connection between them (Ahi and Sefc, 2018). We have recently described such an indirect hierarchical regulatory connection in the pituitary of Atlantic salmon between *vgll3* and *jun*; an upstream regulator of sexual maturation (Ahi et al., 2022a). In mammals, for instance, it is already known that *Six3* and *Six6* together induce the expression of the *Hes5* transcription repressor (Diacou et al., 2018), a negative regulator of neurogenesis which is highly expressed in the anterior part of developing hypothalamus (Aujla et al., 2015). A well-known direct downstream target of *Hes5* in brain is a gene called *Fbw7* or *Fbxw7* (Sancho et al., 2013), which encodes a member of the F-box protein family and controls neural stem cell differentiation in various parts of the brain (Hoeck et al., 2010). The transcriptional repression of *Fbw7* by Hes5 is essential for the correct specification of neural cell fates (Sancho et al., 2013). Interestingly, it has been recently shown that Ythdf2 is a direct substrate for Fbw7 and their interaction leads to proteolytic degradation of Ythdf2 (Xu et al., 2021). These findings in mammals indicate a potential hierarchical regulatory axis consisting of Six6/Hes5/Fbw7/Ythdf2 which can explain a molecular link between Six6 and Ythdf2. However, this model still does not explain the differential transcriptional regulation of *ythdf2* by distinct *six6* genotypes in the hypothalamus of Atlantic salmon.

Another interesting finding in this study was the sex-specific differential expression of *alkbh5-1* with higher expression in the female hypothalamus (Fig. 7). The product of this gene is a well known m^6^A demethylase which acts as an eraser of m^6^A on RNA and involved in a variety of biological processes but has thus far mainly been studied in the context of human diseases (J. Wang et al., 2020). *Alkbh5* deficient male mice, have significantly impaired fertility resulting from apoptosis that affects meiotic metaphase-stage spermatocytes (Zheng et al., 2013). A later study showed the broader role of *Alkbh5* in normal spermatogenesis and male fertility by controlling splicing and stability of long 3′-UTR mRNAs in male germ cells (Tang et al., 2017). During the oogenesis of Xenopus laevis, the expression level of *Alkbh5 was* found to be very high in the growing oocyte (Qi et al., 2016). But knowledge on potential role of *Alkbh5* in female sexual maturation has remained limited. During mammalian neural development, *Alkbh5* is highly expressed in different parts of brain including a dense expression in hypothalamus (Du et al., 2020). Future studies are required to understand potential role of *alkbh5* in hypothalamus in respect to sexual maturation. The higher hypothalamic expression of *alkhb5-1* in the hypothalamus of female Atlantic salmon might indicate an overall lower level of m^6^A methylation in females but the biological relevance of this sex-specific difference in unclear and no prior evidence has been shown such a bias in other species.

A paralog of a reader gene, *ythdf1-3*, showed sex-specific differential expression, i.e. higher expression in the female hypothalamus (Fig. 7). *Ythdf1* is a well-known m^6^A reader which facilitates translation initiation through synergistic interactions with initiation factors and promotion of ribosome loading on m^6^A-modified mRNAs (Wang et al., 2015). Although the specific role(s) of *Ythdf1* has not been explored in the hypothalamus, *Ythdf1* has been shown to play various roles in central nervous system such as hippocampus-dependent learning and memory processing (Shi et al., 2018), spinal axon guidance (Zhuang et al., 2019), cerebellar fiber growth (Yu et al., 2021), and nerve regeneration (Livneh et al., 2020). Interestingly, it is shown in mice that *Ythdf1* is a direct post-transcriptional target for Alkbh5 and its mRNA m^6^A demethylation is mediated by eraser activity of Alkbh5 which leads to increased expression level of *Ythdf1* (Han et al., 2021). This is consistent with our findings, and the identical expression patterns between *alkbh5-1* and *ythdf1-3*, both with higher expression in the hypothalamus of females, suggests a regulatory connection between them in salmon similar to the one observed in mice.

## Conclusions

This study provides the first characterization of expression patterns in all types of m^6^A RNA modification regulator genes, i.e. writers, erasers and readers, in fish hypothalamus. Moreover, it demonstrates the existence of paralog-specific variation in expression of these genes in the hypothalamus of Atlantic salmon indicating potential differences in their function in this organ. On the other hand, both genotype- and maturity status-dependent expression differences were detected for a well-known reader gene involved in mRNA degradation, *ythdf2-2*, i.e. higher expression in immature and *late* (L) puberty allele of *six6* in the hypothalamus of male Atlantic salmon. This suggests for the first time in a vertebrate species a potential role of *ythdf2-2* in sexual maturation in hypothalamus. In addition to *ythdf2-2*, we also found sex-specific expression (females > males) of an eraser gene, *alkbh5-1* and its downstream reader target, *ythdf1-3*, which may implicate an unknown sex-dependent mechanism regulating m^6^A RNA modification regulator genes in the hypothalamus of Atlantic. Further transcriptional and functional investigations are required to understand the underlying regulatory mechanisms linking genotype, maturity status and sex to these m^6^A RNA modification regulators.

## Acknowledgements

We acknowledge Jaakko Erkinaro and staff at the Natural Resources Institute Finland (Luke) hatchery in Laukaa and members of the Evolution, Conservation and Genomics research group for their help in coordinating and collecting gametes for crosses. We thank Nikolai Piavchenko and a number of other assistants for help with fish husbandry and Iikki Donner for dissection and laboratory assistance.

## Author Contributions

EP, MF, CRP conceived the study; EP and MF performed experiments; MH, MF and EP developed methodology and analyzed the data; EP, MF and CRP interpreted results of the experiments; EP, MF and CRP drafted the manuscript, with EP having the main contribution, and all authors approved the final version of manuscript.

## Funding Source Declaration

Funding was provided by Academy of Finland (grant numbers 307593, 302873, 327255 and 342851), and the European Research Council under the European Articles Union’s Horizon 2020 research and innovation program (grant no. 742312).

## Competing financial interests

Authors declare no competing interests

## Ethical approval

Animal experimentation followed European Union Directive 2010/63/EU under license ESAVI/35841/2020 granted by the Animal Experiment Board in Finland (ELLA).

## Data availability

All the gene expression data generated during this study are included in this article as supplementary file.

## Notes

### Competing Interest Statement

The authors have declared no competing interest.

